# Blast Traumatic Brain Injury Induces Long-Term Alterations in Inflammatory Gene Expression in Chinchilla Brains

**DOI:** 10.1101/2024.05.24.595770

**Authors:** Rebecca Schmitt, Kathiravan Kaliyappan, Supriya D. Mahajan, Vijaya Prakash Krishnan Muthaiah

**Affiliations:** Department of Rehabilitation Science, School of Public Health and Health Professions, University at Buffalo, South Campus, Buffalo, NY 14215, USA; Department of Medicine, Jacob School of Medicine and Biomedical Sciences, University at Buffalo, Buffalo, NY 14203, USA

**Keywords:** Neuroinflammation, Chinchilla, Blast Injury, Pro-inflammatory markers

## Abstract

Blast traumatic brain injury (bTBI) due to high-intensity impulsive noise exposure from explosions and munitions exposure is highly prevalent among military personnel, which leads to diffuse brain injury resulting in a spectrum of brain dysfunction and cognitive deficits. The resultant prolonged neuroinflammation and consequent failure of inflammation resolution is a key contributor to long-term complications, including post-traumatic stress disorder and early-onset of neurodegenerative disease; however, there is little evidence for the duration and extent of long term neuroinflammation in distinct brain regions. To investigate this, due to human-like audiogram, we use chinchillas as an *in-vivo* bTBI model to analyze the relative gene expression of inflammatory markers (TNFα, TGFβ2, Gal1, HSP90, S100B, NRGN, MAPK14, IL8, NFL, and BDNF) in the hippocampus, striatum, and higher centers of the auditory pathway 90 days following varying intensities of blast exposures (144 dB, 155 dB, and 172 dB sound pressure level). Our study revealed aberrant gene expression across all analyzed brain regions and all injury conditions; however, no specific pattern emerged. Many of the inflammatory markers were downregulated, suggesting a possible attempt by the brain to overcome prior inflammation. Conversely, the hippocampus, striatum, inferior colliculus, and medial geniculate body all exhibited upregulation of inflammatory markers, including the TBI prognostic marker, S100B. Thus, chinchilla brains exhibit evidence of prolonged neuroinflammation 90 days following injury, even during mild blast exposure. Ultimately, the observed alterations in the gene expression of inflammatory markers may contribute to the long-term neurological dysfunction and neurodegenerative disease experienced by veterans and other bTBI patients.

## 1.0 Introduction

Traumatic brain injury (TBI) is a complex and heterogeneous injury that occurs as a consequence of external force on the brain and results in significant cognitive and physical impairment [1–3]. While TBI is a major cause of death and disability among the general population of the United States [4], military personnel are at an increased risk due to the environmental factors of training and combat [5–8]. Indeed, the Department of Defense (DOD) reports nearly 500,000 TBIs among U.S. service members from 2000 and 2023 [9]. Of these, a majority are obtained as a result of blast overpressure (BOP) from exposure to acoustic shock waves generated by explosives, otherwise known as blast TBI (bTBI) [10–16]. However, despite ongoing research efforts, the pathophysiology of bTBI is incompletely understood.

In contrast to traditional TBIs, which result from acceleration and deceleration forces generated by direct physical impact [17], In bTBIs predominantly primary injury occur due to the compressive and tensile components of BOP [18]. Previous studies indicate that the subsequent neurological effects of bTBI are diffuse, widespread, and variable between different brain regions [18–21]. Injuries within the striatum and hippocampus are closely associated with significant aftermath problems, including cognitive impairment [17,22,23], epilepsy [23,24], post-traumatic stress disorder (PTSD) [1,25], and affective disorder [1,6,8,23], marking them as major regions of interest (ROI) for bTBI research. However, we and others have shown that bTBI also results in auditory neurodegeneration [26–29], which can cause chronic auditory dysfunctions such as hearing loss, tinnitus, and central auditory processing disorders [1,29] which by itself potentiating the early onset of neurodegeneration. Furthermore, bTBI induces chronic vascular remodeling in a range of brain regions, including the cerebellum, hypothalamus, thalamus, neocortex, hippocampus, basal ganglia, piriform cortex, and amygdala, which leads to perivascular astrocytic degeneration [2,19] and degradation of the blood-brain barrier (BBB) [30–33]. This chronic vascular remodeling is associated with an increased risk of post-bTBI stroke or hemorrhage as well as emotional disturbances, such as PTSD [15,19].

A robust neuroinflammatory response is a major feature of TBI and bTBI secondary injury [19,20,34], which refers to delayed tissue, cellular, or molecular changes that are not directly caused by the initial mechanical force [20,35]. Even low-level blast exposure, the mildest form of bTBI that results from outgoing munitions [13], results in increased neuroinflammation as characterized by the release of pro-inflammatory mediators as well as the activation of microglia and astrocytes [19,36,37]. This neuroinflammation is associated with changes in gene expression connected to cell death and metabolism [31,32,37,38], contributing to bTBI-related neurodegeneration [2], auditory dysfunction [26], and BBB degradation [21] leading to failure of inflammation resolution. Notably, chronic unresolved neuroinflammation can lead to increased production of amyloid β and tau proteins, which increases the risk for neurodegenerative diseases such as Alzheimer’s and Parkinson’s disease in bTBI patients [5,15,26]. Nevertheless, there is currently little insight into the duration of neuroinflammation nor the rate of its resolution in distinct brain regions following bTBI.

In our current study, we examine the gene expression of inflammatory mediators associated with neuroinflammation due to bTBI primary (over-pressurizing force) injury in Chinchillas, 90 days following a singular BOP. To understand the long-term changes in these pro-inflammatory mediators throughout the brain, ROIs of the central auditory neuraxis (CAN) are analyzed, including the medial geniculate body (MGB), inferior colliculus (IC), and auditory cortex (AC), as well as other key regions of the brain such as the right and left striatum and hippocampus. Understanding the duration and extent of chronic neuroinflammation associated with primary injury of bTBI may facilitate the discovery of novel therapeutics and prognostic bTBI biomarkers.

## 2.0 Materials and Methods

### In vivo TBI animal Model

Adult male chinchillas of about 350-500g were purchased from Ryerson Chinchilla Ranch, USA, and housed in the Laboratory Animal Facilities, at the University at Buffalo. All experiments were performed according to the regulations and guidelines of the United States Department of Agriculture (USDA) and approved by the Institutional Animal Care and Use Committee (IACUC) of the University at Buffalo. Chinchillas were unilaterally (ipsilateral – left side and contralateral – right side) exposed to a BOP (impulse) with a maximum energy spectrum below 10 kHz, as described below. The animals were grouped into control, 144dB, 155 dB, and 172dB sound pressure level (SPL) groups (measured using a B & K Type 4938 microphone), with a total of n=15 animals/ group.

### Unilateral Blast-overpressure (BOP)

The animals were anesthetized with Xylazine (4mg/kg) and Ketamine (40mg/kg), then exposed to a single impulse blast wave unilaterally inside a soundproof single-walled chamber. In this quasi-experimental blast exposure, both the ipsilateral and contralateral ears were un-occluded, conforming to the standard animal models for blast-induced neurotrauma [39]. The impulse blast wave was generated by a custom-built acoustic shock tube (NIOSH design developed by Mark Cauble Precision Inc, USA) [40], fitted with a compressed airflow system, pressure chamber, and catenoid horn, which can generate impulse noise up to 40 psi (275 kPa) with its energy greatest below several kHz. The pressure chamber was set at 23 psi, 26 psi, and 35 psi to generate BOPs of ∼144 dB, ∼155 dB, and ∼172dB SPL, respectively. The initial acoustic over-pressure occurred in less than 2 milliseconds and the acoustic pressure data of each blast exposure was measured using a B & K ¼ inch pressure field microphone (Type 4938) placed near the tragus of the ipsilateral and contralateral chinchilla.

### RNA Extraction

Tissue RNA was extracted by an acid guanidinium-thiocyanate-phenol-chloroform method, using TRIzol^®^ reagent. 500μl TRIzol^®^ reagent was added to the brain samples weighing 10-30 mg and homogenized in beadBug microtube homogenizer (benchmark Scientific model D1030) for 60 sec at 3500 rpm. 100μl of chloroform was added to each sample and incubated at room temperature for 10 mins, followed by centrifugation at 13000 rpm for 15 mins at 4C. 300 μL of isopropanol was added to each sample, then stored at -80C overnight to precipitate the RNA. The samples were were centrifuged at 13000 rpm for 30 min at 4C and the resulting RNA pellet was washed twice with 1ml of 75% ethanol followed by air drying for 5 mins. RNA was re-constituted with 25μl of DEPC H20, then quantified using a Nano-Drop ND-1000 spectrophotometer. Isolated RNA was stored at -80°C until use.

### RT-qPCR

500 ng of total RNA extracted as outlined above, was used for the First-Strand cDNA Synthesis Kit (GE Healthcare, Piscataway, NJ), according to the manufacturer’s instructions. The resulting cDNA was diluted 1:10 in DEPC H20 and 1 μL was employed as the template in PCR reactions using well-validated PCR primers obtained from the Real-TimePrimers.com. A list of primers and their sequences can be found in Table 1. Primers were provided as a 20 μL solution containing both forward and reverse primers at a final concentration of 10 μM in 10 mM Tris-HCl (pH 7.5) and 0.1 mM EDTA, which were diluted with RNAase/DNase free water as needed before use. The SYBR^®^ Greenmaster mix that contained dNTPs, MgCl2, and DNA polymerase (Bio-Rad, Hercules, CA) was used for gene detection, with a final primer concentration of 0.1 μM. PCR conditions were as follows: 95 °C for 3 min, followed by 40 cycles of 95 °C for 40 s, 60 °C for 30 s, and 72 °C for 1 min; the final extension was at 72 °C for 5 min. Gene expression was calculated using the comparative CT method. The threshold cycle (Ct) of each sample was determined, the relative level of a transcript (2ΔCt) was calculated by obtaining ΔCt (test Ct − β-actin Ct), and the transcript accumulation index (TAI) was calculated as TAI = 2−ΔΔCT.

### Statistical Analysis

Statistical analysis was performed using t-tests between uninjured control and blast-injured groups in GraphPad Prism (GraphPad Software, Inc). P value at the alpha level of < 0.05 was considered statistically significant. <u>A detailed description of the statistical analysis can be found in the supplemental information.</u>

## 3.0 Results

### 3.1 Hippocampus

Although chronic neuroinflammation resulting from bTBI can increase a patient’s risk for cognitive and physical disability as well as neurodegenerative disease, little is known about the duration and extent of long-term neuroinflammation that results from the failure of inflammation resolution. To investigate this, Chinchillas were exposed to a BOP of 144 dB, 155 dB, and 172 dB, generated by a custom-built acoustic shock tube that was fitted with a compressed airflow system, pressure chamber, and catenoid horn; then brain tissues were harvested after 90 days. The relative gene expression of TBI-related inflammatory markers in the brains of injured animals in comparison to sham control was then determined in various brain regions using RT-qPCR. Given that injuries in the hippocampus are associated with many post-bTBI complications, we began with an investigation of the left and right hippocampus. Here, TNFα, HSP90, BDNF, and NFL were all determined to be downregulated by greater than 25% under all injury conditions (Left: **Fig. 1A**, Right: **Fig. 1B**). IL8 was found to be downregulated by at least 40% under all injury conditions in the right hippocampus (**Fig. 1B**) but exhibited no significant difference in the left hippocampus (**Fig. 1A**). In the left hippocampus, Gal1 was downregulated by 79% following 144 dB BOP and showed no significant difference following 155 dB and 172 dB BOP, while MAPK14 exhibited a 99% decrease following both 144 dB and 155 dB BOP but showed no significant difference following 172 dB injury (**Fig. 1A**). In the right hippocampus, TGFβ2 and S100B were downregulated by 95% and 32%, respectively, following 144 dB BOP but showed no significant difference after 155 dB and 172 dB injury (**Fig. 1B**). Thus, the majority of analyzed inflammatory markers were significantly downregulated 90 days after BOP.

**Fig 1.**
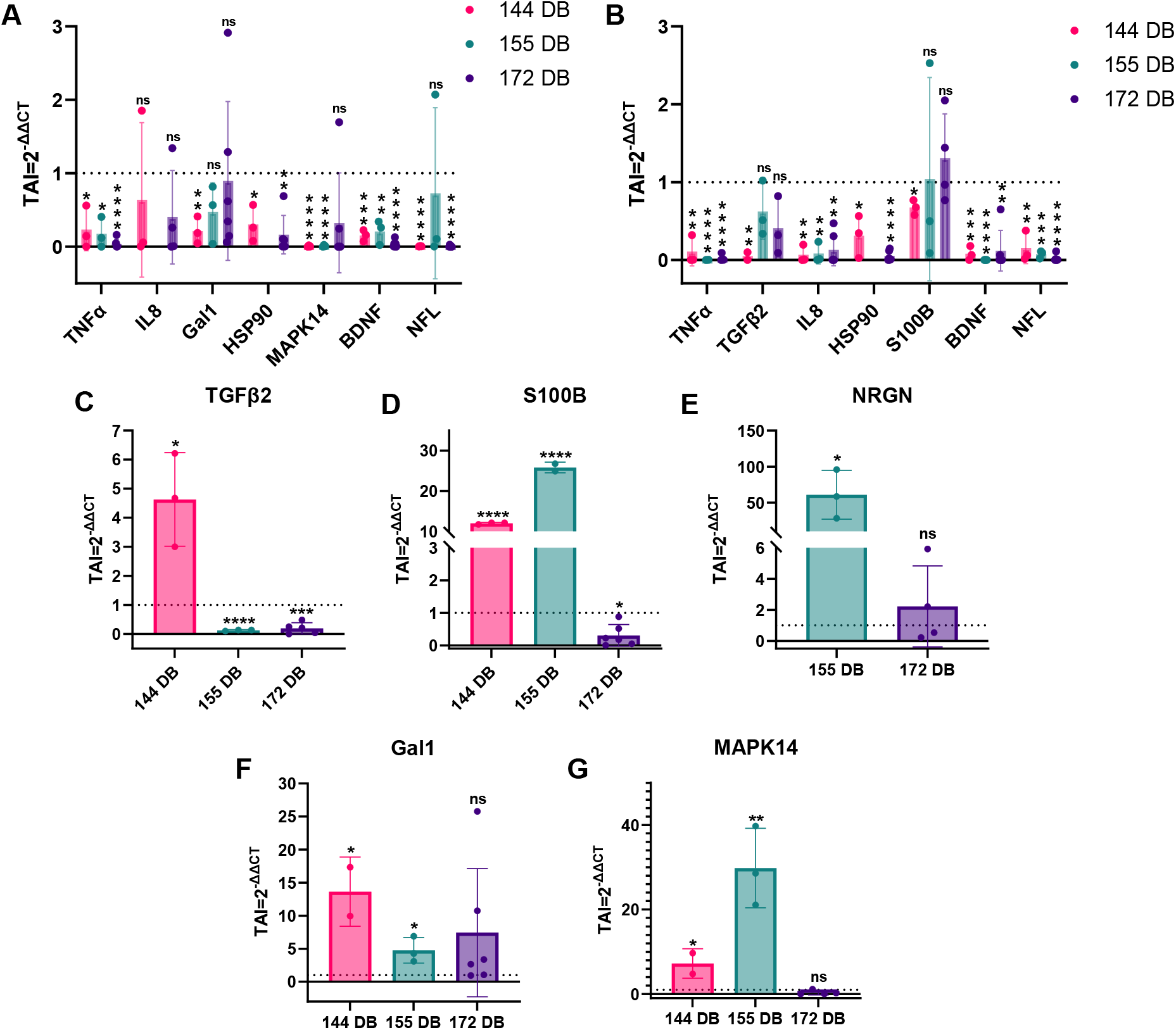
Post-Blast Hippocampal Gene Expression – The relative gene expression of (**A**) TNFα, IL8, Gal1, HSP90, MAPK14, BDNF, and NFL in the left hippocampus as well as (**B**) TNFα, TGFβ2, IL8, HSP90, S100B, BDNF, and NFL in the right hippocampus of Chinchillas following 144 dB, 155 dB, and 172 dB BOP compared to uninjured controls. The relative gene expression of (**C**) TGFβ2, (**D**) S100B, and (**E**) NRGN in the left hippocampus as well as (**F**) Gal1 and (**G**) MAPK14 in the right hippocampus of Chinchillas following 144 dB, 155 dB, and 172 dB BOP.

Several inflammatory markers did, however, show increased expression 90 days following injury in both the left and right hippocampus. In the left hippocampus, TGFβ2 was upregulated 3.6-fold following 144 dB BOP but downregulated by greater than 80% following 155 dB and 172 dB injury (**Fig. 1C**). In the same brain region, S100B was upregulated 11-fold following 144 dB BOP and 25-fold following 155 dB BOP, and downregulated by 69% after 172 dB injury (**Fig. 1D**). 155 dB injury also resulted in a drastic 60-fold upregulation of NRGN, while 172 dB injury caused no significant difference in the left hippocampus (**Fig. 1E**). In the right hippocampus, Gal1 (**Fig. 1F**) and MAPK14 (**Fig. 1G**) were each found to be overexpressed 12.6-fold and 6.2-fold, respectively, following 144 dB BOP as well as 3.7-fold and 29-fold following 155 dB BOP, while 172 dB BOP caused no significant difference.

### 3.2 Striatum

The striatum is another major brain region associated with post-bTBI complications and, thus, was an important ROI in our study. We continued to use our Chinchilla bTBI model in which TNFα, IL8, HSP90and NFL were found to be downregulated by at least 76% in both the left (**Fig. 2A**) and right (**Fig. 2B**) striatum of the brain 90 days following injury, under all conditions. In the left striatum, TGFβ2 and NRGN showed no significant difference following 144 dB and 172 dB injury, while 155 dB BOP resulted in an 83% and 72% decrease in relative gene expression, respectively, compared to uninjured controls (**Fig. 2A**). In the same brain region, S100B demonstrated a 93% decrease after 144 dB BOP and an 83% decrease after 155 dB BOP, while 172 dB injury resulted in no significant difference (**Fig. 2A**). 144 dB and 172 dB also resulted in a 98-99% decrease in MAPK14 in the left striatum, while 155 dB resulted in no significant change (**Fig. 2A**). In the right striatum, MAPK14 and BDNF were found to be downregulated by 96-97% following 144 dB BOP, exhibited no significant difference after 155 dB BOP and were downregulated by 99.9% and 83%, respectively, following 172 dB BOP (**Fig. 2B**). In the same brain region, Gal1 was downregulated by 97% following 144 dB injury and 91% after 155 dB injury but exhibited no change following 172 dB injury (**Fig. 2B**). The only inflammatory-related gene found upregulated in the right striatum was S100B, which was increased by 2-fold and 9-fold following 144 dB and 172 dB, respectively (**Fig. 2C**). However, in the same brain region, 172 dB injury did not cause any significant difference to S100B (**Fig. 2C**).

**Fig 2.**
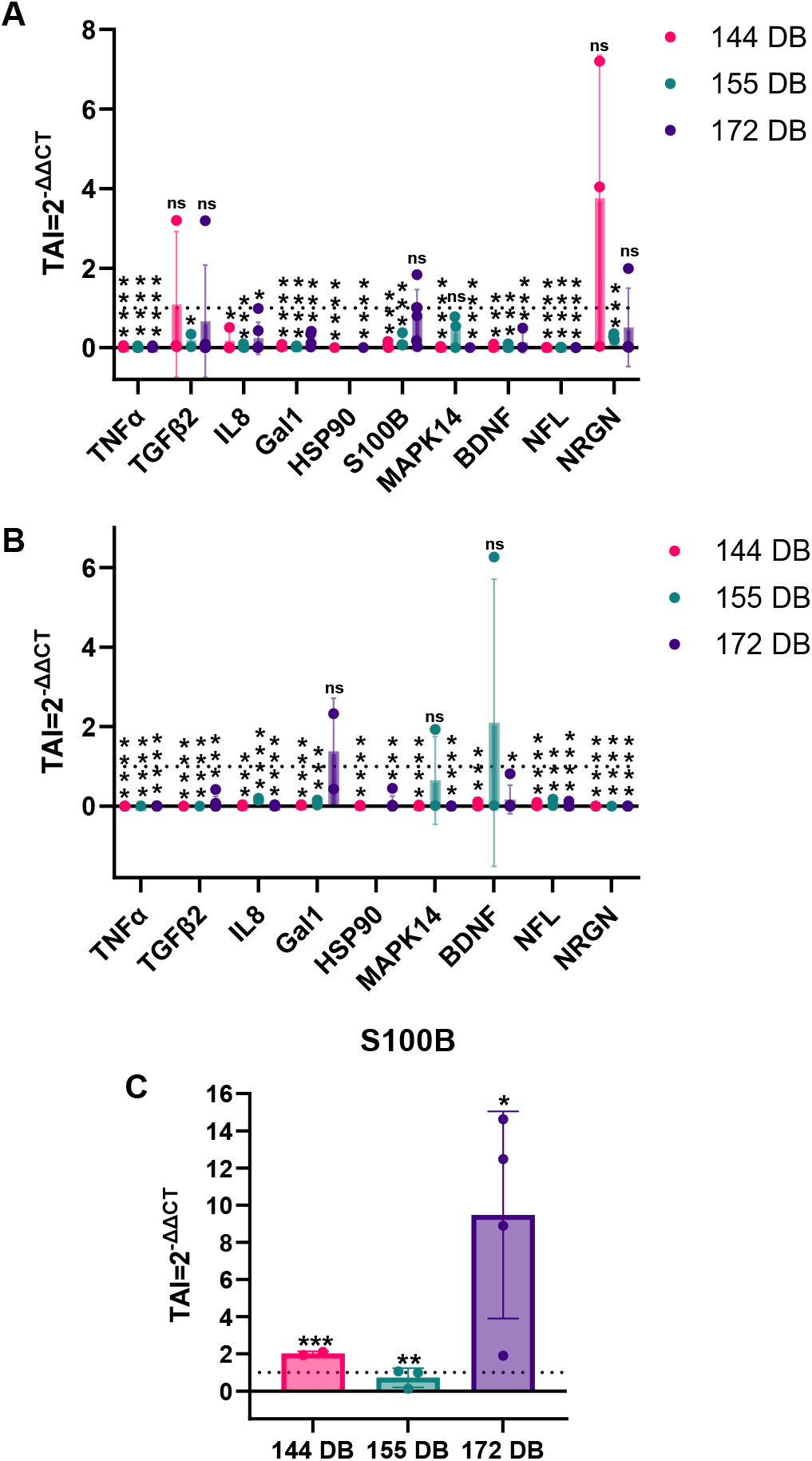
Post-Blast Striatal Gene Expression – The relative gene expression of TNFα, TGFβ2, IL8, Gal1, HSP90, MAPK14, BDNF, and NFL in the (**A**) left striatum and, apart from S100B, (**B**) the right striatum following blast injury at intensities ranging from 144 dB to 172 dB. (**C**) The relative gene expression of S100B in the left striatum.

### 3.3 Central Auditory Neuraxis

#### 3.3.1 Auditory Cortex

Given that we previously identified significant neurodegeneration in the CAN of Chinchilla brains and that auditory dysfunction is a major feature of bTBI, we were particularly interested in determining whether CAN neuroinflammation persisted 90 days following blast injury. In both the left (**Fig. 3A**) and right (**Fig. 3B**) AC, nearly all analyzed genes were found to be significantly downregulated under all injury conditions, 90 days following BOP, in comparison to uninjured controls: including, TNFα, TGFβ2, IL8, HSP90, MAPK14, and NFL. However, Gal1 and BDNF exhibited no significant change in expression following a 172 dB BOP in the left AC, while 144 dB and 155 dB resulted in greater than a 90% decrease of both genes (**Fig. 3A**). In the right AC, Gal1 and NRGN showed no significant differences in gene expression under all analyzed injury conditions (**Fig. 3B**). Furthermore, S100B was found to have no significant difference following 155 dB BOP in the right AC but was downregulated by 56% and 83% following 1144 dB and 72 dB BOP, respectively (**Fig. 3B**). BDNF showed no significant change following 144 dB and 172 dB BOP but was downregulated by 83% in the same brain region (**Fig. 3B**).

**Fig 3.**
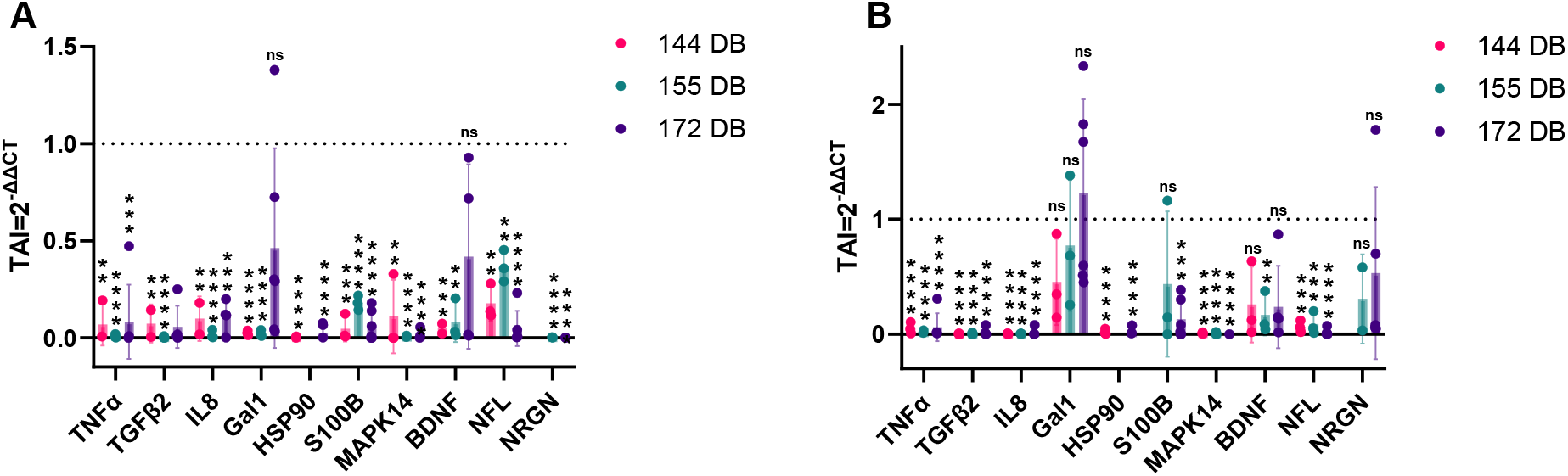
Post-Blast AC Gene Expression – Relative gene expression of TNFα, TGFβ2, IL8, Gal1, HSP90, MAPK14, BDNF, and NFL in (**A**) left and (**B**) right AC following 144 dB, 155 dB, and 172 dB BOP.

#### 3.3.2 Inferior Colliculus

In the IC, TGFβ2 and BDNF were both found to be downregulated by more than 80% under all injury conditions. TNFα was also found to be downregulated by greater than 85% following 155 dB and 172 dB BOP but showed no significant difference after 144 dB injury (**Fig. 4A**). IL8 showed no significant difference after 144 dB and 155 dB injury but was downregulated by 96% after 172 BOP (**Fig. 4A**). MAPK14, on the other hand, was downregulated by 99% after 144 dB and 155 dB injury and was not found to have any significant difference following 172 dB BOP (**Fig. 4A**). HSP90 was also found to be downregulated by 70% after 144 dB BOP and showed no significant difference after 172 dB BOP (**Fig. 4A**). S100 (**Fig. 4B**) and NRGN (**Fig. 4D**) were both found to be upregulated by 12-fold and 62-fold respectively following 144 dB BOP, while S100B (**Fig. 4B**) and Gal1 (**Fig. 4C**) were upregulated by 20-fold and 17-fold following 155 dB BOP. However, S100B was found to be downregulated by 69% following 172 dB BOP (**Fig. 4B**). Gal1 was also found to be downregulated by 69% following 144 dB BOP and displayed no significant difference following 172 dB BOP (**Fig. 4C**). NRGN showed no significant difference at either 155 dB BOP or 172 dB BOP (**Fig. 4D**).

**Fig 4.**
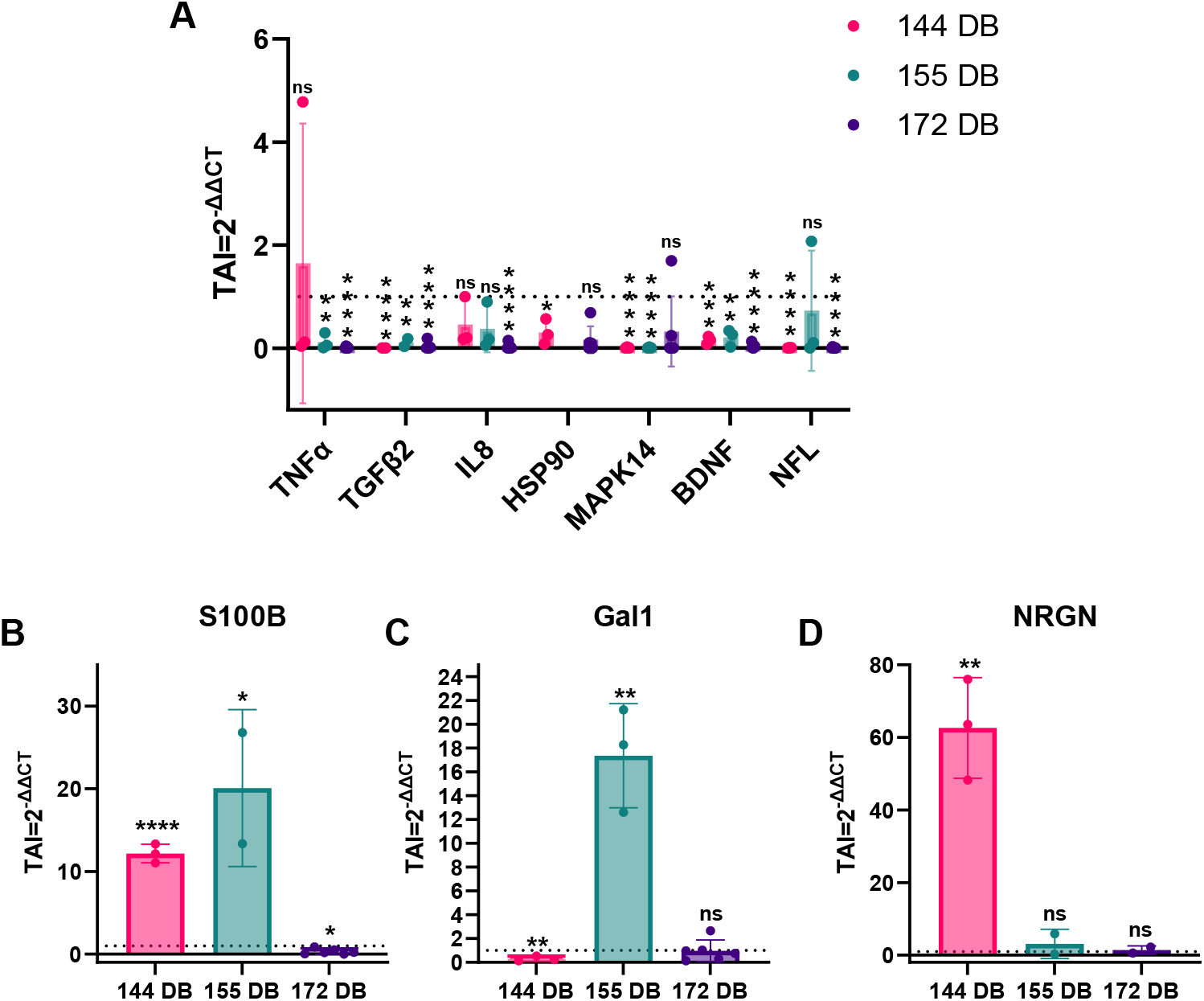
Post-Blast IC Gene Expression – (**A**) Relative gene expression of TNFα, TGFβ2, IL8, HSP90, MAPK14, BDNF, and NFL in the right IC following 144 dB, 155 dB, and 172 dB BOP. Relative gene expression of (**B**) S100B, (**C**) Gal1, and (**D**) NRGN in the right IC under the same injury conditions.

#### 3.3.3 Medial Geniculate Body

In the MGB, TNFα and MAPK14 were both found to be significantly downregulated by > 90% under all injury conditions, while there was no significant difference found between uninjured controls and each injury condition for HSP90 and NFL (**Fig. 5A**). In this brain region, IL8 was increased by 6.7-fold following 144 dB injury but no significant difference was found at 155 dB and 172 dB (**Fig. 5B**). TGFβ2, on the other hand, was drastically overexpressed by 74-fold to 274-fold across all injury conditions (**Fig. 5C**). Although no statistically significant difference was found in the expression of Gal1 under any injury condition, Gal1 did trend towards overexpression by 5.0-fold following 172 dB injury (**Fig. 5D**). BDNF exhibited a 7.8-fold overexpression following 172 dB injury as well but showed no significant difference following 144 dB or 155 dB BOP (**Fig. 5E**).

**Fig 5.**
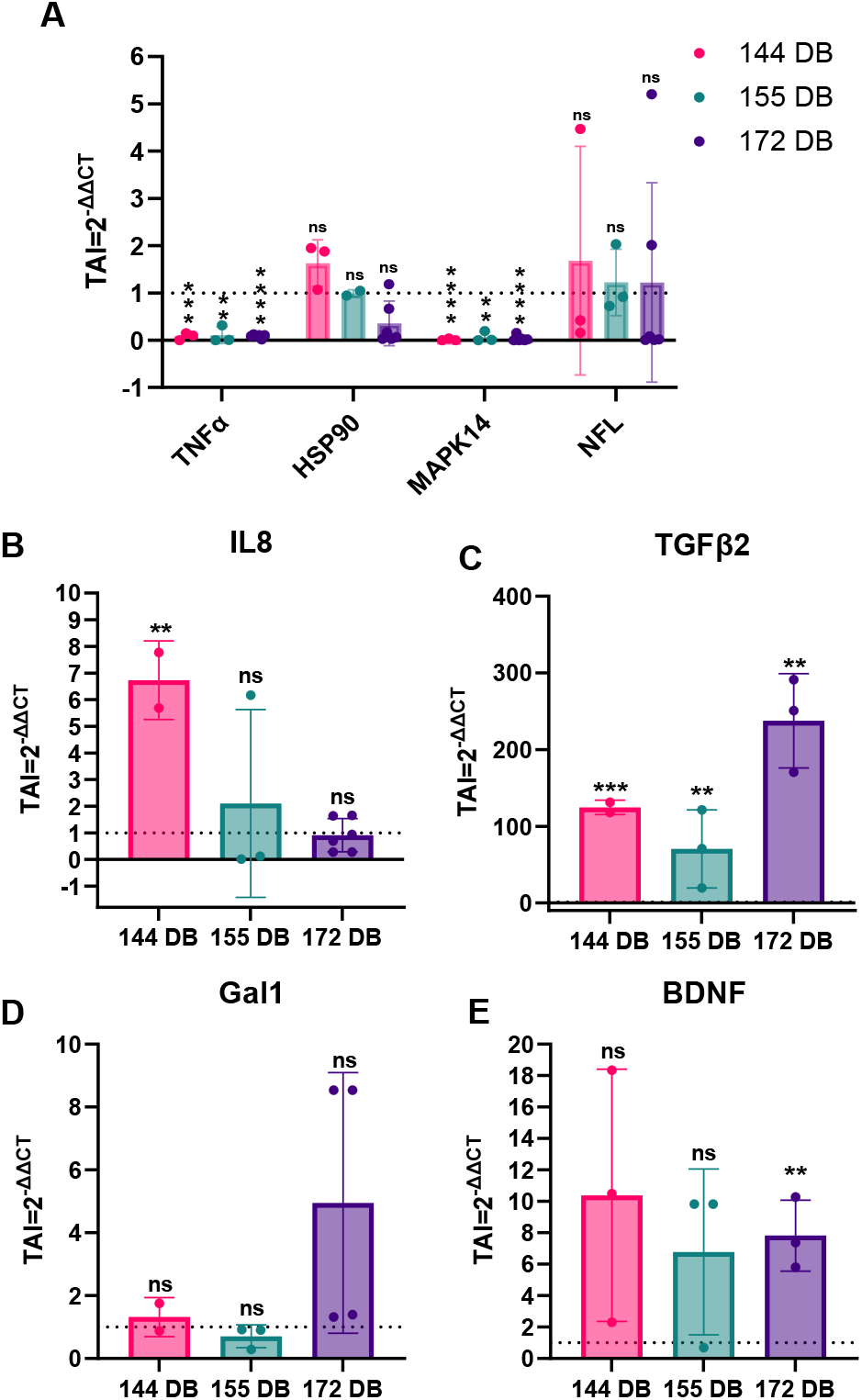
Post-Blast MGB Gene Expression – (**A**) Relative gene expression of TNFα, HSP90, MAPK14, and NFL in the right MGB following 144 dB, 155 dB, and 172 dB BOP. Relative gene expression of (**B**) IL8, (**C**) TGFβ2, (**D**) Gal1, and (**E**) BDNF in the right MGB under the same injury conditions.

To better understand how the inflammatory proteins encoded by our analyzed genes interact with each other and other potential players in bTBI, we conducted interaction mapping using a gene function prediction program (**Fig. 6**). While weak interactions were identified between most of our analyzed genes, TNFα exhibited a large number of significant interactions with genes both known to be involved in TBI pathophysiology, including NEFL (aka NFL), and those that are lesser studied, such as LIN54, which is critical in cell cycle regulation [41]; SPOCK1, which is found to be downregulated in the cerebrospinal fluid after TBI and is involved in BBB integrity [42,43]; and ATP2A2, which codes for an ATPase that regulates intracellular calcium ion concentrations [44]. Other strong interactions with TNFα include MRPS33, PTPRZ1, SEC16A, SLU7, ERICH2, ST13, NT5DC3, and CARS1, all of which are understudied for their role in TBI. The TBI prognostic marker, S100B, also showed an indirect interaction with TNFα through NKAPD1, which enables identical protein binding [45]. Additional interactions with S100B include SPG21, which is involved in suppressing T-cell activation, as well as BCL2L2, which codes for an anti-apoptotic BCL2-like protein 2 [46]. Aside from interactions with TNFα and S100B, significant interactions were found between the neuroprotective BDNF and AGO3, which is involved in RNA silencing and post-transcriptional gene regulation [47], as well as between HSP90 and CERS2, which is related to cell growth and death [48]. Ultimately, many major interactions were found between the genes analyzed in this study and potential genes of interest for bTBI pathogenesis.

**Fig 6.**
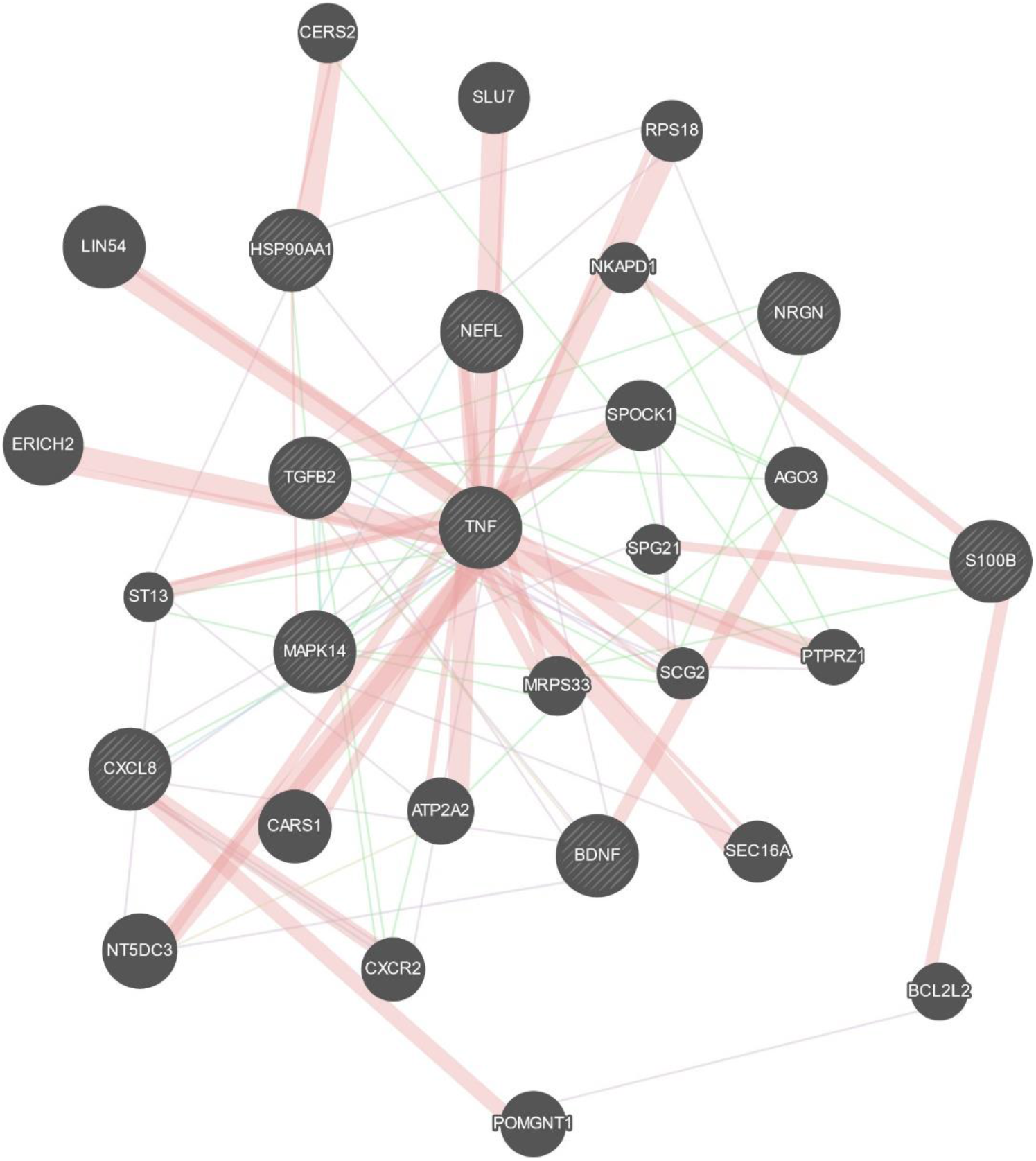
Schematic representation of the interaction between inflammatory markers. CERS2 – Ceramide Synthase; SLU7 - SLU7 homolog, splicing factor; RPS18 – Ribosomal protein S18 LIN54 - lin-54 DREAM MuvB core complex component; HSP90AA1 – Heat Shock Protein alpha family class A member 1; NKAPD1 - NKAP domain containing 1; NEFL - neurofilament light (NFL); ERICH2 -glutamate-rich 2; ST13 - ST13 Hsp70 interacting protein; MAPK14 - mitogen-activated protein kinase 14; MRPS33 – Mitochondrial Ribosomal Protein S33; SCG2 -Secretogranin II; SEC16A - SEC16 homolog A, endoplasmic reticulum export factor; BDNF-Brain-derived Neurotropic Factor; S100B - S100 calcium-binding protein B; NT5DC3 - 5’-nucleotidase domain containing 3; TNF – Tumor Necrosis Factor; TGF – Transforming Growth Factor; SPG21 - SPG21 abhydrolase domain containing, maspardin; PTPRZ1 - Protein tyrosine phosphatase receptor type Z1; CXCL8 - C-X-C motif chemokine ligand 8; CXCR2 - C-X-C motif chemokine receptor 2; AGO3 - argonaute RISC catalytic component 3; NRGN – Neurogranin; SPOCK1 - SPARC (osteonectin), cwcv and kazal like domains proteoglycan 1; POMGNT1 - protein O-linked mannose N-acetylglucosaminyltransferase 1

## 4.0 Discussion

A study by Veteran Affairs indicates that even a mild bTBI is associated with post-traumatic stress disorder as well as depressive and neurobehavioral symptoms in correlation with the blast exposure severity [1]. These psychological disturbances are attributed to diffuse brain injury resulting from the compressive and tensile components of BOP as well as subsequent secondary effects [18,20,35,38]. Indeed, the impact of BOP can be observed throughout the brain, with widespread neurodegeneration [2,26], vascular remodeling [2,19], and auditory dysfunction [28,29] as major bTBI features. Using Chinchillas, whose audiogram is similar to humans, as an *in vivo* bTBI model, we previously found that blast exposure increases degenerating neurons and apoptotic cells at the auditory pathway, especially at the cochlear nucleus, IC, and AC [27]. This, in part, could be due to the over-activation of the DNA repair enzyme, Poly ADP Ribose Polymerase-1 (PARP-1), in response to BOP which can be recuperated by a known PARP-1 inhibitor, 3-aminobenzamide, as we’ve shown in mouse auditory cells (HEI-OC1) [49]. Additionally, in an earlier *in-vitro* study using human brain microvascular cells, we demonstrated that blast exposure disrupted the integrity of the BBB which was evidenced by the downregulation in the gene expression of various tight junction proteins such as Claudin 1,2 and 3, ZO1, and JAM2 as well as blast-induced apoptosis [31,32]. Overall, blast exposure results in dysregulated gene network activity related to cell death and metabolism [31,37,38].

The damage to various regions of the brain resulting from BOP often triggers neuroinflammation across the cortical and sub-cortical centers [19,34]. A failure of this inflammation to resolve is related to chronic neurological manifestations following blast injury in Veterans, including increased risk for neurodegenerative diseases such as Alzheimer’s and Parkinson’s [5,26,50]. Nevertheless, few evidence is available on the status of neuroinflammation post-blast injury, particularly with respect to duration or rate of inflammation resolution in various brain regions. Considering the blast-induced neurological sequelae, in this study, we investigated the gene expression of inflammatory markers (TNFα, TGFβ, Gal1, S100B, NRGN, MAPK14, IL8, and BDNF) in various ROIs within the brains of Chinchillas 90 days after unilateral blast exposure of various intensities (144-, 155- and 172-dB SPL). Most of the neurological impairments associated with memory, learning, and executive function are associated with striatal and hippocampal dysfunction [51], thus the striatum and hippocampus serve as major ROIs in this study. However, as mentioned above, BOP leads to significant auditory neurodegeneration, hence we also focused on higher centers of auditory pathways such as the AC, IC, and MGB.

Our results indicate that many inflammatory markers were drastically downregulated in the hippocampus (**Fig. 1A** and **1B**) and striatum (**Fig. 2A** and **2B**) 90 days following blast exposure to each injury condition, including TNFα, HSP90, and BDNF, with several other markers exhibiting no significant difference. This suggests a potential attempt by the brain to reduce neuroinflammation in the long term, resulting in overcompensation concerning the downregulated genes. Importantly, aberrant downregulation of these inflammatory markers is still dysregulation and may contribute to long-term neurological dysfunction in bTBI patients. The downregulation of TNFα is particularly intriguing given that our gene function interaction mapping revealed several significant interactions between TNFα and genes encoding other regulatory proteins, including SPOCK1 (Testican-1) and ATP2A2 (SERCA2), which have not been substantially investigated in connection with bTBI (**Fig. 6**). Notably, loss of function of ATP2A2 can lead to a risk for psychiatric disorders, including depression, bipolar disorder, and schizophrenia [44]. Further investigation of the downregulation of inflammatory markers as well as the genes and proteins they interact with in the hippocampus and striatum could provide valuable insight into the long-term deficits experienced because of bTBI.

Despite the downregulation of many inflammatory markers in the hippocampus and striatum, several genes were found to be upregulated under certain conditions. The upregulation of TGFβ2 observed in the left hippocampus following 144 dB BOP (**Fig. 1C**) supports an attempt to correct bTBI neuroinflammatory effects, as it directs microglia to an anti-inflammatory activation state [52,53]. However, the upregulation of the TBI biomarker, NRGN [54], in the left hippocampus (**Fig. 1E**), inflammatory markers Gal1 (**Fig. 1F**) and MAPK14 (**Fig. 1G**) in the right hippocampus, and the TBI prognostic marker S100B [55] in the left hippocampus (**Fig. 1D**) and striatum (**Fig. 2C**) under various blast intensities, all suggest continued upregulation of pro-inflammatory pathways under each respective injury condition. Furthermore, our gene map analysis revealed a significant interaction between S100B and the anti-apoptotic gene BCL2l2 (BCL2-like protein 2) (**Fig. 6**) [46]. While anti-apoptotic pathways will reduce neuronal death following TBI, it is possible that upregulation of such a pathway could allow dysregulated cells to perpetuate, further contributing to neural dysfunction.

Any level of blast exposure leads to long-term degeneration of the auditory pathway and results in chronic auditory dysfunctions such as hearing loss, tinnitus, central auditory processing disorders, and poor speech intelligibility [29,56]. Interestingly, evidence is evolving that tinnitus can trigger cognitive impairment and early-onset dementia [57,58], which could be affiliated with unresolved neuroinflammation. In our study, all analyzed pro-inflammatory markers were reduced or found to have no significant difference in both the right and left AC irrespective of the severity of the exposure (**Fig. 3A** and **3B**), again suggesting an overcompensation for initial neuroinflammation. This overcompensation for neuroinflammation can also be seen by the downregulation of many genes in the right IC (**Fig. 4A**) and MGB (**Fig. 4C**) as well as the overexpression of TGFβ2 (**Fig. 5C**) under all injury conditions and the overexpression of BDNF (**Fig. 5E**), which supports neuronal repair [59], following 172 dB BOP in the right MGB. However, in the right IC, we found the TBI biomarkers S100B (**Fig. 4B**) and NRGN (**Fig. 4D**) were overexpressed following 144 dB BOP, with S100B and the inflammatory marker Gal1 (**Fig. 4C**) also being overexpressed after 155 dB injury, suggesting continued neuroinflammation in this ROI. Furthermore, in the right MGB, the pro-inflammatory markers IL8 (**Fig. 5B**) and Gal1 (**Fig. 5D**) were both overexpressed following 172 dB BOP, with IL8 also being upregulated following 144 dB injury (**Fig. 5B**). Thus, these findings suggest a continued activation of inflammatory pathways in the right IC and MGB of the CAN 90 days following injury.

In conclusion, we observed significant alterations in gene expression for TNFα, TGFβ2, Gal1, HSP90, S100B, NRGN, MAPK14, IL8, NFL, and BDNF in multiple ROIs of the Chinchilla brain, including the hippocampus, striatum, and higher centers of the CAN, 90 days following blast exposure of varying intensities. The extent and nature of the changes observed varied between brain regions and blast intensities; however, no specific trend or pattern emerged. In most instances, the analyzed genes were found to be significantly downregulated, suggesting a potential attempt by the brain to counteract prior inflammation but resulting in overcompensation. This concept was further substantiated by an upregulation of anti-inflammatory or neuroprotective genes in the hippocampus and the MGB. However, there were several pro-inflammatory markers overexpressed in the hippocampus, striatum, right IC, and right MGB, including the TBI prognostic marker S100B, thus indicating a possibility of persistent unresolved neuroinflammation 90 days following injury. The gene dysregulation specifically observed in the CAN demonstrates that continued auditory dysfunction following bTBI is complex and requires further investigation. All in all, the dysregulation of our analyzed genes could contribute to the long-term neurological dysfunction experienced by Veterans and other bTBI patients as well as potentially contribute to the onset of neurodegenerative disease.

## Acknowledgements

We would like to acknowledge Satyabrata Parida, PhD and Professor Robert Burkard for their help with Matlab support and acoustic calibrations respectively.

## Conflict of Interest

The authors whose names are listed in the title page certify that they have NO affiliations with or involvement in any organization or entity with any financial interest (such as honoraria; educational grants; participation in speakers’ bureaus; membership, employment, consultancies, stock ownership, or other equity interest; and expert testimony or patent-licensing arrangements), or non-financial interest (such as personal or professional relationships, affiliations, knowledge or beliefs) in the subject matter or materials discussed in this manuscript. The contributing authors do not have any financial interest that affects the outcome of the study.

## Credit Distribution

Vijaya Prakash Krishnan Muthaiah and Supriya Mahajan conceptualized the study design, mobilized resources, established the blast, sample harvest, and interpretation, manuscript writing. Rebecca Schmidt supported gene expression experiments, data collection, statistical analysis. Kathiravan Kaliyappan contributed with blast exposures and sample collections.

## Notes

Conflict of Interest: None.

### Competing Interest Statement

The authors have declared no competing interest.

