## Supplemental Information for "Blast Traumatic Brain Injury Induces Long-Term Alterations in Inflammatory Gene Expression in Chinchilla Brains"

|  |  | 144 dB | | | 155 dB | | | 172 dB | | |
| --- | --- | --- | --- | --- | --- | --- | --- | --- | --- | --- |
|  | **Gene** | **t ratio** | **df** | **q value** | **t ratio** | **df** | **q value** | **t ratio** | **df** | **q value** |
| **Left** | TNFα | 3.532 | 3 | 0.025978 | 5.325 | 3 | 0.017426 | 20.37 | 6 | <0.000001 |
|  | TGFβ2 | 3.913 | 4 | 0.0174* | 65.38 | 4 | <0.0001* | 6.946 | 6 | 0.0004* |
|  | IL8 | 0.4629 | 3 | 0.389551 |  |  |  | 1.251 | 4 | 0.14091 |
|  | Gal1 | 5.744 | 3 | 0.010585 | 1.781 | 3 | 0.174754 | 0.1293 | 6 | 0.390139 |
|  | HSP90 | 3.786 | 3 | 0.025978 |  |  |  | 4.292 | 6 | 0.003891 |
|  | S100B | 83.01 | 4 | <0.0001* | 35.29 | 3 | <0.0001* | 3.481 | 7 | 0.0103* |
|  | MAPK14 | 128.3 | 3 | 0.000002 | 113.9 | 3 | 0.000006 | 1.336 | 6 | 0.139411 |
|  | BDNF | 14.76 | 3 | 0.000909 | 6.402 | 3 | 0.015596 | 26.41 | 6 | <0.000001 |
|  | NFL | 588.5 | 3 | <0.000001 | 0.3147 | 3 | 0.625065 | 121.4 | 6 | <0.000001 |
|  | NRGN |  |  |  | 3.062 | 4 | 0.0376* | 0.7918 | 5 | 0.4644* |
| **Right** | TNFα | 6.513 | 3 | 0.007427 | 752.7 | 3 | <0.000001 | 36.04 | 6 | <0.000001 |
|  | TGFβ2 | 18.88 | 2 | 0.00376 | 1.416 | 3 | 0.101671 | 2.121 | 3 | 0.041752 |
|  | IL8 | 11.33 | 3 | 0.002976 | 9.29 | 3 | 0.001333 | 5.657 | 6 | 0.000662 |
|  | Gal1 | 4.576 | 3 | 0.0196* | 3.35 | 4 | 0.0286* | 1.11 | 7 | 0.3039* |
|  | HSP90 | 3.409 | 3 | 0.024342 |  |  |  | 19.6 | 6 | <0.000001 |
|  | MAPK14 | 3.382 | 3 | 0.0430* | 5.302 | 4 | 0.0061* | 2.024 | 5 | 0.0988* |
|  | S100B | 4.587 | 3 | 0.013105 | 0.03962 | 3 | 0.326864 | 0.7164 | 4 | 0.148133 |
|  | BDNF | 13.43 | 3 | 0.002976 | 1465 | 3 | <0.000001 | 4.52 | 6 | 0.001623 |
|  | NFL | 5.749 | 3 | 0.008446 | 26.37 | 3 | 0.000081 | 27.23 | 5 | <0.000001 |
|  | *indicates p-value instead of q-value | | | | | | | | | |

**Supporting Information**

**Contents**

Pg 1 Hippocampal statistical summary

Pg 2 Striatal statistical summary

Pg 3 Auditory Cortex statistical summary

Pg 4 Inferior Colliculus and Medical Geniculate Body statistical summary

**Table S1** Summary of the statistical analysis of the relative gene expression of inflammatory markers in the left and right hippocampus of Chinchillas, 90 days after exposure to blast injury of varying intensities (144 dB, 155 dB, and 172 dB).

**Table S2** Summary of the statistical analysis of the relative gene expression of inflammatory markers in the left and right striatum of Chinchillas, 90 days after exposure to blast injury of varying intensities (144 dB, 155 dB, and 172 dB).

|  |  | 144 dB | | | 155 dB | | | 172 dB | | |
| --- | --- | --- | --- | --- | --- | --- | --- | --- | --- | --- |
|  | **Gene** | **t ratio** | **df** | **q value** | **t ratio** | **df** | **q value** | **t ratio** | **df** | **q value** |
| **Left** | TNFα | 41.65 | 3 | 0.000023 | 53.22 | 3 | 0.000015 | 74.71 | 6 | <0.000001 |
|  | TGFβ2 | 0.06733 | 3 | 0.288019 | 5.191 | 2 | 0.013318 | 0.3165 | 5 | 0.386026 |
|  | IL8 | 3.779 | 3 | 0.0123 | 24.56 | 3 | 0.000101 | 2.531 | 6 | 0.032192 |
|  | Gal1 | 31.28 | 3 | 0.000044 | 89.25 | 3 | 0.000005 | 5.906 | 5 | 0.002001 |
|  | HSP90 | 180.7 | 3 | 0.000001 |  |  |  | 188.3 | 6 | <0.000001 |
|  | S100B | 15.25 | 3 | 0.000265 | 6.341 | 3 | 0.003434 | 0.3891 | 6 | 0.386026 |
|  | MAPK14 | 84.06 | 3 | 0.000006 | 1.872 | 3 | 0.053148 | 681.3 | 6 | <0.000001 |
|  | BDNF | 25.79 | 3 | 0.000065 | 23.58 | 3 | 0.000101 | 4.717 | 4 | 0.007739 |
|  | NFL | 240.9 | 2 | 0.000017 | 355.1 | 3 | <0.000001 | 715.8 | 6 | <0.000001 |
|  | NRGN | 1.03 | 3 | 0.127482 | 12.94 | 3 | 0.000503 | 0.6598 | 4 | 0.344314 |
| **Right** | TNFα | 5116 | 3 | <0.000001 | 383.2 | 3 | <0.000001 | 366.6 | 6 | <0.000001 |
|  | TGFβ2 | 9522 | 2 | <0.000001 | 1708 | 3 | <0.000001 | 7.427 | 6 | 0.000103 |
|  | IL8 | 59.24 | 2 | 0.000288 | 28.19 | 3 | 0.00005 | 85.96 | 4 | <0.000001 |
|  | Gal1 | 222.4 | 3 | <0.000001 | 19.16 | 3 | 0.000125 | 0.3981 | 2 | 0.163627 |
|  | HSP90 | 134.8 | 2 | 0.000083 |  |  |  | 6.899 | 6 | 0.000132 |
|  | S100B | 16.37 | 3 | 0.0005* | 0.9391 | 4 | 0.4008* | 2.572 | 5 | 0.0499* |
|  | MAPK14 | 83.18 | 3 | 0.000007 | 0.4251 | 3 | 0.179367 | 1313 | 6 | <0.000001 |
|  | BDNF | 21.06 | 3 | 0.000266 | 0.4084 | 3 | 0.179367 | 3.084 | 5 | 0.006905 |
|  | NFL | 30.05 | 3 | 0.000105 | 13.92 | 3 | 0.00027 | 26.46 | 6 | <0.000001 |
|  | NRGN | 1.52E+09 | 2 | <0.000001 | 11286681 | 2 | <0.000001 | 10185650 | 4 | <0.000001 |
|  | *indicates p-value instead of q-value | | | | | | | | | |

**Table S3** Summary of the statistical analysis of the relative gene expression of inflammatory markers in the left and right auditory cortex of Chinchillas, 90 days after exposure to blast injury of varying intensities (144 dB, 155 dB, and 172 dB).

|  |  | 144 dB | | | 155 dB | | | 172 dB | | |
| --- | --- | --- | --- | --- | --- | --- | --- | --- | --- | --- |
|  | **Gene** | **t ratio** | **df** | **q value** | **t ratio** | **df** | **q value** | **t ratio** | **df** | **q value** |
| **Left** | TNFα | 11.54 | 3 | 0.002119 | 133.1 | 3 | 0.000002 | 6.446 | 6 | 0.000167 |
|  | TGFβ2 | 13.29 | 2 | 0.00729 | 345.2 | 3 | <0.000001 | 11.63 | 5 | 0.000028 |
|  | IL8 | 11.09 | 2 | 0.008159 | 60.59 | 3 | 0.000015 | 14.55 | 4 | 0.000037 |
|  | Gal1 | 112.4 | 3 | 0.000007 | 85.2 | 3 | 0.000006 | 1.401 | 6 | 0.042555 |
|  | HSP90 | 359.7 | 3 | <0.000001 |  |  |  | 35.75 | 6 | <0.000001 |
|  | S100B | 19.36 | 3 | 0.000912 | 29.55 | 3 | 0.00011 | 16.16 | 6 | 0.000002 |
|  | MAPK14 | 6.3 | 3 | 0.008159 | 357.2 | 3 | <0.000001 | 61.3 | 6 | <0.000001 |
|  | BDNF | 37.58 | 2 | 0.001607 | 11.71 | 3 | 0.001519 | 1.631 | 4 | 0.039992 |
|  | NFL | 12.32 | 3 | 0.002095 | 10.43 | 3 | 0.001902 | 14.08 | 6 | 0.000003 |
|  | NRGN |  |  |  | 1731 | 2 | 0.000001 | 1419 | 3 | <0.000001 |
| **Right** | TNFα | 85.76 | 3 | 0.000004 | 10.34 | 6 | 0.000024 | 25.66 | 3 | 0.000046 |
|  | TGFβ2 | 173 | 3 | <0.000001 | 43.51 | 6 | <0.000001 | 480.1 | 3 | <0.000001 |
|  | IL8 | 618.9 | 3 | <0.000001 | 42.1 | 6 | <0.000001 | 436.5 | 3 | <0.000001 |
|  | Gal1 | 0.5365 | 3 | 0.282286 | 0.3839 | 6 | 0.216422 | 1.947 | 3 | 0.037048 |
|  | HSP90 |  |  |  | 44.53 | 6 | <0.000001 | 56.91 | 3 | 0.000006 |
|  | S100B | 1.195 | 3 | 0.160642 | 6.915 | 6 | 0.000196 |  |  |  |
|  | MAPK14 | 137.9 | 3 | 0.000001 | 2208 | 6 | <0.000001 | 237.4 | 3 | <0.000001 |
|  | BDNF | 6.173 | 3 | 0.005763 | 2.852 | 5 | 0.013541 | 3.019 | 3 | 0.01639 |
|  | NFL | 12.27 | 3 | 0.000942 | 45.99 | 6 | <0.000001 | 25.16 | 3 | 0.000046 |
|  | NRGN | 2.525 | 2 | 0.073613 | 0.834 | 5 | 0.148903 |  |  |  |

**Table S4** Summary of the statistical analysis of the relative gene expression of inflammatory markers in the right inferior colliculus of Chinchillas, 90 days after exposure to blast injury of varying intensities (144 dB, 155 dB, and 172 dB).

|  | 144 dB | | | 155 dB | | | 172 dB | | |
| --- | --- | --- | --- | --- | --- | --- | --- | --- | --- |
| **Gene** | **t ratio** | **df** | **q value** | **t ratio** | **df** | **q value** | **t ratio** | **df** | **q value** |
| TNFα | 0.3189 | 3 | 0.333599 | 7.537 | 3 | 0.009748 | 91.22 | 6 | <0.000001 |
| TGFβ2 | 5499 | 2 | <0.000001 | 11.83 | 2 | 0.009748 | 17.48 | 6 | <0.000001 |
| IL8 | 1.555 | 3 | 0.109936 | 1.839 | 3 | 0.164901 | 20.85 | 5 | <0.000001 |
| Gal1 | 5.937 | 4 | 0.004** | 6.474 | 4 | 0.0029* | 0.04658 | 7 | 0.9642* |
| HSP90 | 3.786 | 3 | 0.019587 |  |  |  | 4.292 | 6 | 0.000865 |
| S100B | 17.06 | 4 | <0.0001* | 3.811 | 3 | 0.0318* | 3.481 | 7 | 0.0103* |
| MAPK14 | 128.3 | 3 | 0.000001 | 113.9 | 3 | 0.000008 | 1.336 | 6 | 0.033193 |
| BDNF | 14.76 | 3 | 0.000511 | 6.402 | 3 | 0.009748 | 26.41 | 6 | <0.000001 |
| NFL | 588.5 | 3 | <0.000001 | 0.3147 | 3 | 0.65111 | 121.4 | 6 | <0.000001 |
| NRGN | 5.954 | 3 | 0.0095* | 0.7274 | 2 | 0.5426* | 0.4507 | 2 | 0.6963* |
| *indicates p-value instead of q-value | | | | | | | | | |

**Table S5** Summary of the statistical analysis of the relative gene expression of inflammatory markers in the right medical geniculate body of Chinchillas, 90 days after exposure to blast injury of varying intensities (144 dB, 155 dB, and 172 dB).

|  | 144 dB | | | 155 dB | | | 172 dB | | |
| --- | --- | --- | --- | --- | --- | --- | --- | --- | --- |
| **Gene** | **t ratio** | **df** | **q value** | **t ratio** | **df** | **q value** | **t ratio** | **df** | **q value** |
| TNFα | 16.69 | 3 | 0.000473 | 6.671 | 3 | 0.010405 | 33.7 | 6 | <0.000001 |
| TGFβ2 | 24.88 | 3 | 0.0001* | 2.362 | 4 | 0.0775* | 6.659 | 4 | 0.0026* |
| IL8 | 7.375 | 3 | 0.0052* | 0.5411 | 4 | 0.6172* | 0.2382 | 7 | 0.8186* |
| Gal1 | 0.944 | 3 | 0.4148* | 1.361 | 4 | 0.2450* | 1.611 | 5 | 0.1682* |
| HSP90 | 1.724 | 3 | 0.123367 | 0.1056 | 2 | 0.701087 | 1.824 | 6 | 0.079414 |
| MAPK14 | 66.64 | 3 | 0.000015 | 11.53 | 3 | 0.00424 | 22.12 | 6 | <0.000001 |
| BDNF | 2.026 | 4 | 0.1127* | 1.896 | 4 | 0.1308* | 5.214 | 4 | 0.0065* |
| NFL | 0.3782 | 3 | 0.368874 | 0.4272 | 3 | 0.701087 | 0.1406 | 6 | 0.450874 |
| *indicates p-value instead of q-value | | | | | | | | | |
